# A high-density carbon fiber neural recording array technology

**DOI:** 10.1101/320937

**Authors:** Travis L Massey, Samantha R Santacruz, Jason F Hou, Kristofer SJ Pister, Jose M Carmena, Michel M Maharbiz

## Abstract

*Objective:* Microwire and Utah-style neural recording arrays are the predominant devices used for cortical neural recording, but the implanted electrodes cause a significant adverse biological response and suffer from well-studied performance degradation. Recent work has demonstrated that carbon fiber electrodes do not elicit this same adverse response, but these existing designs are not practically scalable to hundreds or thousands of recording sites. We present technology that overcomes these issues while additionally providing fine electrode pitch for spatial oversampling.

*Approach:* We present a 32-channel carbon fiber monofilament-based intracortical neural recording array fabricated through a combination of bulk silicon microfabrication processing and microassembly. This device represents the first truly two-dimensional carbon fiber neural recording array. The density, channel count, and size scale of this array are enabled by an out-of-plane microassembly technique in which individual fibers are inserted through metallized and isotropically conductive adhesive-filled holes in an oxide-passivated microfabricated silicon substrate.

*Main results:* Five-micron diameter fibers are spaced at a pitch of 38 microns, four times denser than state of the art one-dimensional arrays. The fine diameter of the carbon fibers affords both minimal cross-section and nearly three orders of magnitude greater lateral compliance than standard tungsten microwires. Typical 1 kHz impedances are on the order of hundreds of kiloohms, and successful *in vivo* recording is demonstrated in the motor cortex of a rat. 22 total units are recorded on 20 channels, with unit SNR ranging from 0.85 to 4.2.

*Significance:* This is the highest density microwire-style electrode array to date, and this fabrication technique is scalable to a larger number of electrodes and allows for the potential future integration of microelectronics. Large-scale carbon fiber neural recording arrays are a promising technology for reducing the inflammatory response and increasing the information density, particularly in neural recording applications where microwire arrays are already used.

## 1. Introduction

Electrophysiological neural recording is an indispensable and nearly ubiquitous technique in efforts to better understand the nervous system and treat neurological conditions, and correspondingly great efforts are being made to develop arrays that meet the needs of researchers and clinicians for recording from either the central or the peripheral nervous system. Each application has its own specific requirements, but the ideal neural recording array, broadly speaking, satisfies four primary criteria. First, it has a large number of recording sites. Second, those recording sites distribute in the recording volume of interest to detect all relevant single-unit activity, while still spatially oversampling to enable approximate localization of each unit [1–4]. Third, the array elicits the minimal adverse biological response, retarding the recording degradation associated with encapsulation and glial recruitment [5–9]. Fourth, the material stack of the implanted array should last for a significant fraction of a subject’s lifetime; this is primarily a function of the chemical stability of the materials and their interfaces [10–14]. While many existing technologies demonstrate clear progress in each of these areas, existing arrays target at most two of these four criteria [15–17]. Notably, there is a lack of technologies with the requisite full-volume sampling and single-unit detection capabilities that also minimize tissue damage and the initiation of the immune response, particularly during insertion as vasculature is damaged and the blood-brain barrier is compromised [18, 19].

Carbon fiber recording electrodes have shown promise as recording electrodes and have the qualities necessary to form ideal recording electrodes [20–22]. Carbon fiber monofilaments are finer than commercially available microwires, ranging from 4-7 µm in diameter. This is below the threshold for encapsulation [17, 23–26], suggesting carbon fiber’s potential for chronic recording with minimal degradation due to the biological response. Further, graphitized carbon is inert to the biological environment yet electrically conductive [27]. The material has a Young’s modulus of 200-280 GPa for polyacrylonitrile (PAN)-based fibers and as high as 455 GPa for the less common pitch-based fibers [21], aiding insertion, yet the fine diameter affords orders of magnitude more compliance than traditional 25 µm tungsten microwires traditionally found in microwire arrays. Of further benefit compared to traditional microwire materials, carbon fiber is elastic until it yields; Kozai *et. al* demonstrated carbon fiber’s high fracture stress by bending a 6.8 µm carbon fiber to a radius of 500 µm [20]. Despite the many benefits of carbon fiber recording electrodes, the scalability and electrode density requirements remain unsolved. Prior carbon fiber neural recording arrays have been limited by both large (≥150 µm) electrode pitch and the ability to form only an *N* × 2 array [22, 24], or have more closely resembled polytrodes with a cross-section above the encapsulation threshold and an exposed recording area too large to reliably detect high-SNR single units [21].

We propose a carbon fiber microwire-style neural recording array that begins to address these density and scalability criteria by arraying ultrafine 4.8-5.4 µm carbon fibers at a pitch of 38 µm to deliver full-volume sampling while minimizing the adverse biological impact. This work demonstrates a 32-electrode array, but the design is inherently scalable to a larger number of electrodes due to a microfabricated silicon substrate and routing plane through which an arbitrary number of carbon fiber monofilaments may be threaded. Further, key steps of the assembly process can be automated; this is requisite for any neural recording array requiring serial assembly to be truly scalable [28]. The result is the highest-density microwire-style neural recording array with the finest diameter implanted electrodes of any microwire array to date, and a clear path toward scalability to hundreds of recording electrodes. In this work we describe the microfabrication process of silicon substrate as well as the assembly procedure to integrate the carbon fiber microwires with the silicon substrate, followed by a demonstration of the device’s viability for recording from the central nervous system (CNS).

## 2. Materials and methods

### 2.1. Substrate microfabrication process

The microfabrication process flow for the silicon substrate is summarized in Figure 1. First, 150 mm diameter, 280 µm thick double-side-polished prime-grade silicon wafers are cleaned in a 120 °C piranha bath for 10 min, rinsed thoroughly in 18 MΩ water, and spun dry. A layer of 1.05 µm thermal silicon dioxide is grown by wet oxidation in a Tylan furnace at 1050 °C and atmospheric pressure for 3 h. The pattern for the holes extending through the silicon is defined by standard photolithography using a GCA 8500 wafer stepper to expose a 2.8 µm thick film of positive i-line photoresist (OiR 906-12, Fujifilm, Valhalla, NY). Because the 280 µm wafer is thinner than the focal limits of the wafer stepper will tolerate, the process wafer is temporarily bonded during exposure to a 400 µm double-side-polished handle wafer using a single droplet of deionized water in the center of the wafer. Following exposure, the wafers are carefully separated and the photoresist is puddle developed normally in a TMAH-based developer solution (OPD 4262, Fujifilm, Valhalla, NY). Additional photoresist is applied manually to cover any defects in the film. The photoresist is hard baked in a 120 °C oven for 6 h to fully cross-link the photoresist, improving the selectivity of the subsequent plasma etch processes.

**Figure 1.**
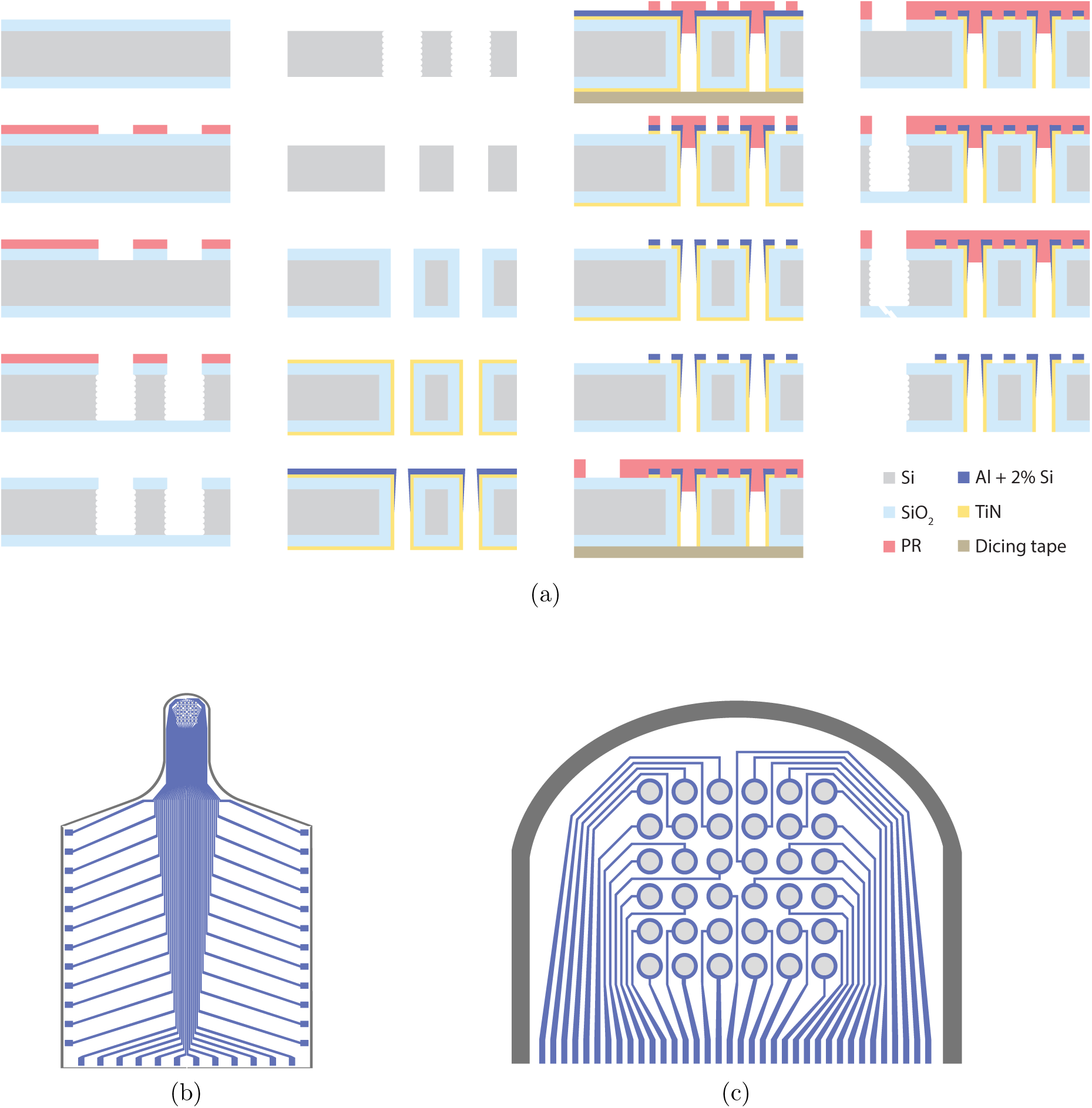
(a) Cross-sectional diagrams of the bulk silicon microfabrication process of the array substrates, as described in Section 2.1. A legend of key materials is provided. (b) Layout (top) view of silicon substrate. (c) Close-up of head of silicon substrate. Through-holes are indicated in light gray, metal traces in blue, and the device outline singulation etch in dark gray.

Etching the holes through the silicon proceeds in two steps. First, the thermal oxide is etched in an inductively coupled plasma (ICP) for 4 min (15 sccm C_4_F_8_, 8 sccm H_2_, 174 sccm He, 1500 W coil, 350 W bias at 13.56 MHz, 0 °C platen, SPTS Technologies Advanced Planar Source Oxide Etch System, Orbotech, Billerica, MA). Next, 20 µm diameter holes are etched through bulk silicon by deep reactive ion etching (DRIE), stopping on the backside thermal oxide (SPTS Technologies ICP-SR Deep Reactive Ion Etch System, Orbotech, Billerica, MA). Detailed etch parameters are provided in the supplementary information (Table S1). Following this etch, the photoresist is stripped in a Matrix 106 Resist Removal System flowing reactive oxygen species (450 s, 250 °C, 400 W, 3.75 Torr O_2_), and the wafers are further cleaned for 10 min in 75 °C RCA2 (5:1:1 H_2_O:HCl:H_2_O_2_) to remove any residual organic residue followed by a rinse in 18 MΩ water as before. The SiO_2_ is etched completely in 60 min in 5:1 buffered hydrofluoric acid (BHF), and the wafers are again rinsed and spun dry. The etched silicon is annealed for 10 min at 1090 °C in 10 Torr pure H_2_ to smooth the scallops resulting from the DRIE process and round the corners at the mouth of each hole (Epi 200 Centura, Applied Materials, Santa Clara, CA) [29].

The wafers are then oxidized as before, and 300 cycles of conductive titanium nitride (TiN) is deposited by plasma-enhanced atomic layer deposition (PEALD) to coat all surfaces of the wafer (Fiji Advanced Atomic Layer Deposition System, Cambridge NanoTech/Veeco Instruments, Waltham, MA), notably including inside the holes, using a modified version of the recipe provided in Burke *et. al* [30]. Following TiN PEALD, the wafer is dehydrated at 200 °C on a hot plate, and 120 nm Al/Si (98%/2%) is deposited in an MRC/TES-944 5 kHz pulsed-DC sputter system (8 mTorr Ar process pressure, 1 kW power, 20 passes at 70 cm/minute; Technical Engineering Services, Santa Cruz, CA). The metallized wafers are lithographically patterned as before with the pattern for the upcoming metal etch, with two notable deviations. First, spinning photoresist now requires that the wafer be backed with a layer of dicing tape (Ultron Systems, 1005R) in order to hold vacuum on the spin chuck due to the through-holes etched previously. This dicing tape is removed during exposure and the post-exposure bake step, and it is reapplied during puddle development, which also occurs on a spin chuck. The second notable deviation is that this layer of photoresist is spun manually, as opposed to using an automated coat track, to accommodate customization of the spin recipe. This customization is necessary due to the severe topography now present on the surface of the wafer and the aggressive feature sizes targeted and proximity to the through-holes. Specifically, the photoresist is dynamically dispensed at 100 RPM for 10 s, after which the spinner is ramped up to 1000 RPM at 100 RPM/s and held at that speed for 30 s before returning to a stop at a rate of −100 RPM/s. Once photolithography is complete, the photoresist is hard baked as before.

Prior to etching the metallization stack, wafers are bonded under vacuum with polyphenyl ether (Santovac 5, SantoLubes LLC, Spartanburg, SC) at 120 °C to standard prime grade handle wafers using a custom wafer bonding tool (Figure S2). The Al and TiN are etched for 50 s in a transformer coupled plasma (TCP) metal etcher (Lam Research Corporation, Fremont, CA) using a 200 W plasma of 90 sccm Cl_2_ and 45 sccm BCl_3_ at 100 W bias. Integrated endpoint detection indicates when the etch has reached the underlying oxide by spectroscopy of the etch products. The process wafer is debonded from the handle wafer and the photoresist stripped by soaking in acetone, with razors inserted around perimeter of the wafer after two hours to provide a small deflection encouraging acetone to flow into the interface and for the wafers to separate. An argon ion mill removes residual TiN from the backside of the wafer (500 V, 300 mA, 10 min, 0° angle, 15 RPM stage rotation; Pi Scientific, Livermore, CA). The high degree of anisotropy inherent to ion milling ensures that the TiN inside the holes is not affected, and less than than 80 nm of SiO_2_ is milled from the backside of the wafer as a consequence of the TiN removal.

Lithographic patterning of the device outlines is performed using a 12 µm film of AZ P4620 photoresist (MicroChemicals GmbH, Ulm, Germany) spun at 2000 RPM and rehydrated, exposed, and developed according to manufacturer specifications. After hard baking the photoresist at 90 °C for 30 min, the SiO_2_ is etched from within the patterned trench in eight 30 s cycles using the SPTS Oxide Etch System described previously. A 90 s cooldown between etch cycles reduces the heating of the photoresist and thus the risk of destructive crack formation. The Si is subsequently etched as before, but the trench etch is carefully timed to stop on the backside SiO_2_ film. The devices are easily removed by breaking the SiO_2_ membrane, and the photoresist is stripped in Microposit Remover 1165 (Dow Chemical Company, Midland, MI), followed by subsequent rinses in acetone, isopropanol (IPA), and deionized (DI) water.

### 2.2. Array assembly process

The process of assembling the complete arrays from the microfabricated substrates begins by epoxying a device substrate to a polyimide flex printed circuit board (PCB) with low-viscosity epoxy (353ND, Epoxy Technology, Inc., Billerica, MA), curing at 150 °C for 1 h. The PCBs and substrates are cleaned in IPA and flowing DI water and dried with nitrogen immediately before wire bonding the substrate to the PCB using a WestBond 747677E wedge bonder outfitted with 25 µm aluminum bond wire and corresponding wedge (WestBond, Anaheim, CA). The wire bonds are protected with the same 353ND epoxy, which is cured as above. A strip of aluminum is taped against the connector solder pads on the back end of the device to discharge all electrostatic charge from the substrate holes and carbon fibers during assembly.

The device and an alignment substrate (see supplementary information for microfabrication details) are mounted on micropositioners, with the silicon parallel to the work surface and the holes of the substrate overhanging the leading edge of each micropositioner. A droplet of water-based silver nanoparticle ink (Novacentrix HPS-030LV, Austin, TX) is applied to the head of the device substrate, covering all thirty-six holes. A doctor blade (durometer 90A polyurethane rubber, McMaster-Carr, Elmhurst, IL) is passed over the surface to force the silver ink into the holes and remove excess from the surface. A second layer of silver epoxy (Atom Adhesives AA-DUCT 24, Fort Lauderdale, FL) is applied to the holes and cleared with a doctor blade in the same way. Excess silver ink is cleaned from the back with the doctor blade, and a small piece of Kapton tape is applied behind the holes as a temporary backstop for the ink, epoxy, and carbon fibers.

The holes of the alignment substrate are precisely aligned above the device substrate using the micropositioners, leaving minimal or no gap between the two substrates, and the 5.2 µm carbon fibers (HexTow IM7, Hexcel, Stamford, CT) are serially threaded through each hole using a third micropositioner. The fibers are temporarily adhered to the probe tip of the third micropositioner by a thin film of cured silicone. Each fiber is lowered into its target hole, and the adhesion between the ink/epoxy and the fiber overcomes the adhesion of the fiber to the silicone, allowing the fiber to remain in the hole when the silicone-coated probe tip is removed. Once all thirty-two fibers are threaded this way (the back-end connector has only thirty-two channels, so four of the thirty-six holes need not be threaded), the alignment device is raised nearly to the tips of the carbon fibers using the micropositioner, ensuring that the fibers are parallel and vertical. The ink and the epoxy are cured in this position in a box oven at 230 °C for 3 h. Following cure, the alignment substrate, backside Kapton tape, and aluminum strip are all removed.

An Omnetics nanostrip connector (NPD-36-VV-GS, Omnetics Connector Corp., Minneapolis, MN) is soldered to the tail of the PCB, and the base is encapsulated in 353ND epoxy as before. An uninsulated 76 µm silver wire is soldered to a reference terminal, and the wire is tacked to the polyimide using UV-curable epoxy (Loctite 3526, Henkel, D¨usseldorf, Germany) to provide strain relief at the solder joint. A 6 mm cube of polystyrene is cut and epoxied on the underside of the PCB beneath the device substrate, as visualized in Figure 3b. The PCB’s tail is then folded back and epoxied to the opposite side of the polystyrene block, and the connector and silver ground wire are sealed and masked with Kapton tape. The entire device is electrically insulated in a conformal 0.8 µm film of parylene-C (Labcoter 2, Specialty Coating Systems, Indianapolis, IN), deposited per manufacturer parameters. Deposition is performed with the device sitting by three contact points on a 100 mm silicon wafer for thickness characterization via ellipsometry. The deposited film is verified free of pinhole defects by placing the wafer in 24% potassium hydroxide (KOH) and monitoring for the evolution of bubbles indicative of KOH reaching and reacting with the silicon surface anywhere other than the device contact points.

To expose the recording sites at the tips of the carbon fibers, the entire device is embedded in a block of Tissue-Tek 4583 embedding compound for cryotoming (Sakura Finetek USA, Torrance, CA) and frozen to −80 °C. The embedded device is mounted into the cryotome held at −55 °C and progressively shaved in 10 µm sections with a TiN-coated blade (C.L. Sturkey, Inc., Lebanon, PA) until the tips of all fibers are exposed, as illustrated in Figure 2a. The embedding compound is thawed and thoroughly rinsed in deionized water. With the tips of the fibers in 1x phosphate buffered saline (PBS), we apply −18 V versus a platinum wire counterelectrode to reduce the impedance through the silver ink, as discussed in Section 4.3. Finally, the recording sites are electroplated with PEDOT:PSS at 7 nA for 60 s using a freshly prepared solution of 0.01 M 3,4-Ethylenedioxythiophene (EDOT) and 0.01 M Poly(sodium 4-styrenesulfonate) (PSS) with molecular weight of approximately 70 kDa (Sigma-Aldrich, St. Louis, MO). A summary of this complete assembly procedure is provided in Figure 2.

**Figure 2.**
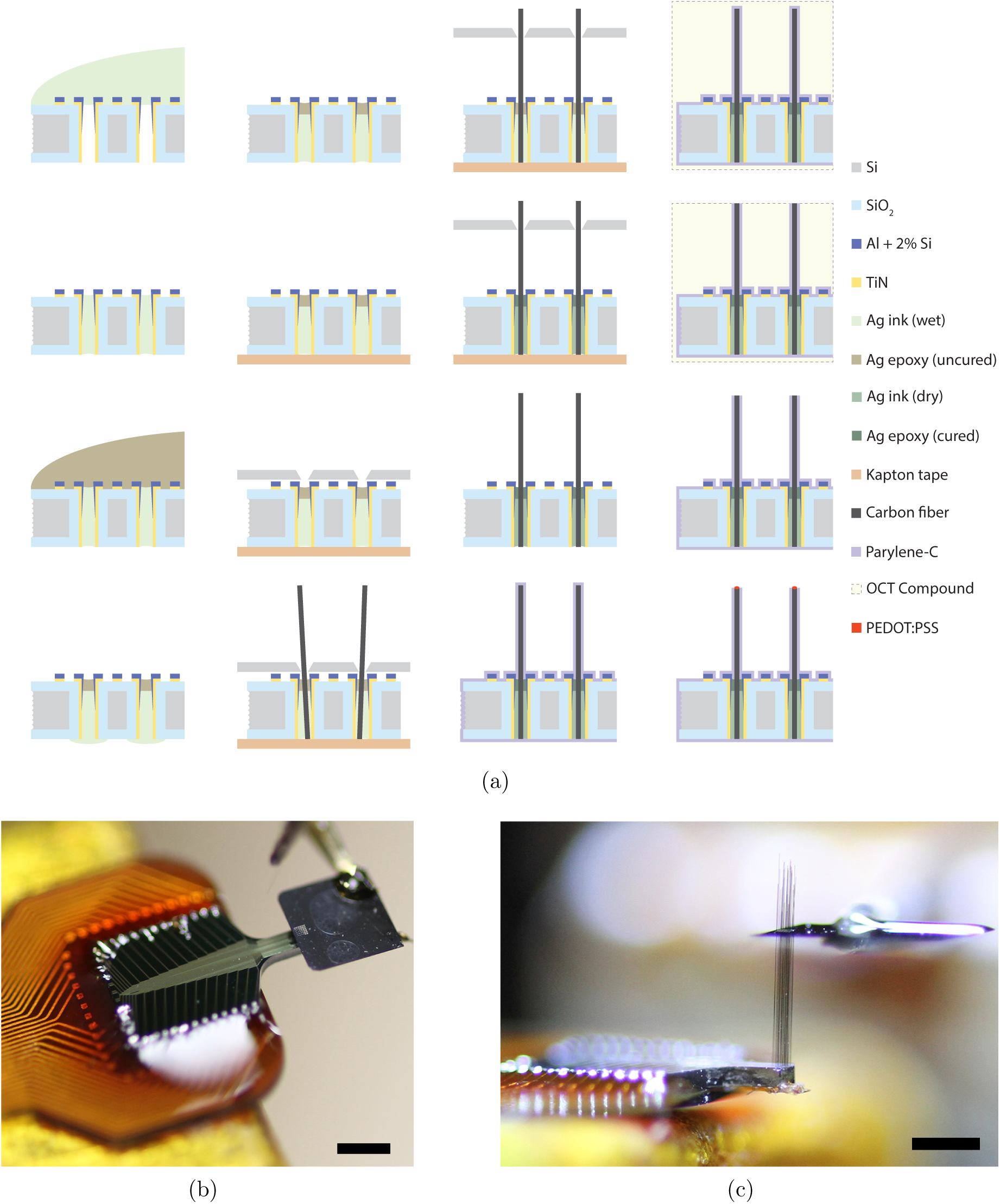
(a) Cross-sectional diagrams of the carbon fiber array assembly process per Section 2.2. A legend of key materials is provided. (b) Photograph of the threading process, with alignment substrate aligned to device substrate and a third probe tip guiding a fiber to the first hole. (c) Photograph of a threaded device during assembly. The alignment substrate has been separated from the device substrate to parallelize the 2.5 mm fibers. Scale bars: 1 mm.

### 2.3. Electrical characterization

All electrical impedance measurements are performed with the recording sites and silver wire reference electrode (if applicable) submerged in PBS. Electrochemical impedance spectroscopy (EIS) is performed over a range of 115 frequencies ranging from 5 Hz to 5 kHz using a nanoZ (White Matter LLC, Seattle, WA) and averaged over 40 cycles. Impedance between every pair of electrodes is measured using a Keysight E4980L precision LCR meter (Santa Rosa, CA) in conjunction with a custom software-controlled multiplexer from [31], taking for each pair the mean of five samples. These pairwise measurements were conducted first in air (open-circuit) before PBS for purposes of identifying potential shorts and quantifying crosstalk between channels. Fibers were submerged in liquefied Field’s metal (51% In, 32.5% Bi, 16.5% Sn; melting point 62 °C) rather than PBS for short-circuit testing (Sections 3.1 and 4.3).

Noise was measured on two recording systems. The first set of noise measurements was performed inside a Faraday cage to minimize electromagnetic interference (EMI) using a PCIE-16AI64SSC-64-B General Standards Corporation (Huntsville, AL) data aquisition card sampling at 20 kHz and FA64I Multi-Channel Systems (Reutlingen, Germany) amplifier with a fifth-order bandpass from 0.1-6000 Hz. The second system was the Plexon Multichannel Acquisition Processor (MAP) recording system sampling at 40 kHz (spike band) and 1 kHz (field potential band) with J2 headstage and PBX-517 preamplifier (Plexon Inc., Dallas, Tx) used for *in vivo* experiments. While lacking a Faraday cage and subject to increased EMI, this latter set of measurements better imitates the noise conditions during *in vivo* testing. Signals were allowed a settling time of 7*τ* before data were recorded, and high-pass filtering data in the first set at 300 Hz (7th-order Type II Chebychev, 80 dB rejection) removed any residual low-frequency drift as well as 60 Hz interference and its significant harmonics with only a 5% reduction in measurement bandwidth. Noise amplitude is calculated as the root mean square of the signal. Data from the second set were filtered only by the built-in preamplifier filters, 500-8800 Hz (field potential band) and 3-200 Hz (slow band). The signal-to-noise ratio (SNR) is calculated as the square of the ratio of the root mean square (RMS) voltage of the mean spike waveform on a given channel to the RMS noise voltage.

### 2.4. Insertion tests

The effect of fiber length and angle on the success of penetration was tested by inserting assembled arrays into 0.6 w/w% agar gel to mimic many mechanical properties of the brain [32–34]. Devices were rotated relative to the agar in 0.2° increments to a maximum of 4.5° using a rotational micropositioner, and were subsequently advanced into the agar such that the direction of motion was off-axis from the fiber by the specified angle. Lengths of 1.4, 1.9, 2.3, 2.8, and 3.5 mm were tested. The agar was shifted slightly between each test so as not to penetrate the same point repeatedly.

### 2.5. In vivo recordings

All experiments were done in accordance with the Animal Care and Use Committee at the University of California, Berkeley, and the National Institutes of Health guidelines. One male Long-Evans rat was acutely implanted unilaterally in M1 with the carbon fiber array under anesthesia. The subject was anesthetized with isoflurane gas throughout the procedure and given dexamethasone approximately 30 min before performing the craniotomy. Single- and multi-unit activity, and local field potentials, were simultaneously recorded with a Plexon MAP. Unit activity was sampled at a rate of 40 kHz. Activity was sorted online and sorting templates were further refined using offline sorting software (OfflineSorter, Plexon Inc., Dallas TX). Post-hoc analysis of neural data was performed using custom written software (Python).

## 3. Results

### 3.1. Recording arrays can be fabricated and assembled

Fabricated substrates and completed devices are shown in Figure 3. The devices shown were assembled with 0.5-1 mm-long carbon fibers for ease of imaging, but typical implanted devices had 2.5-3 mm fibers, such as that shown in Figure 2c.

**Figure 3.**
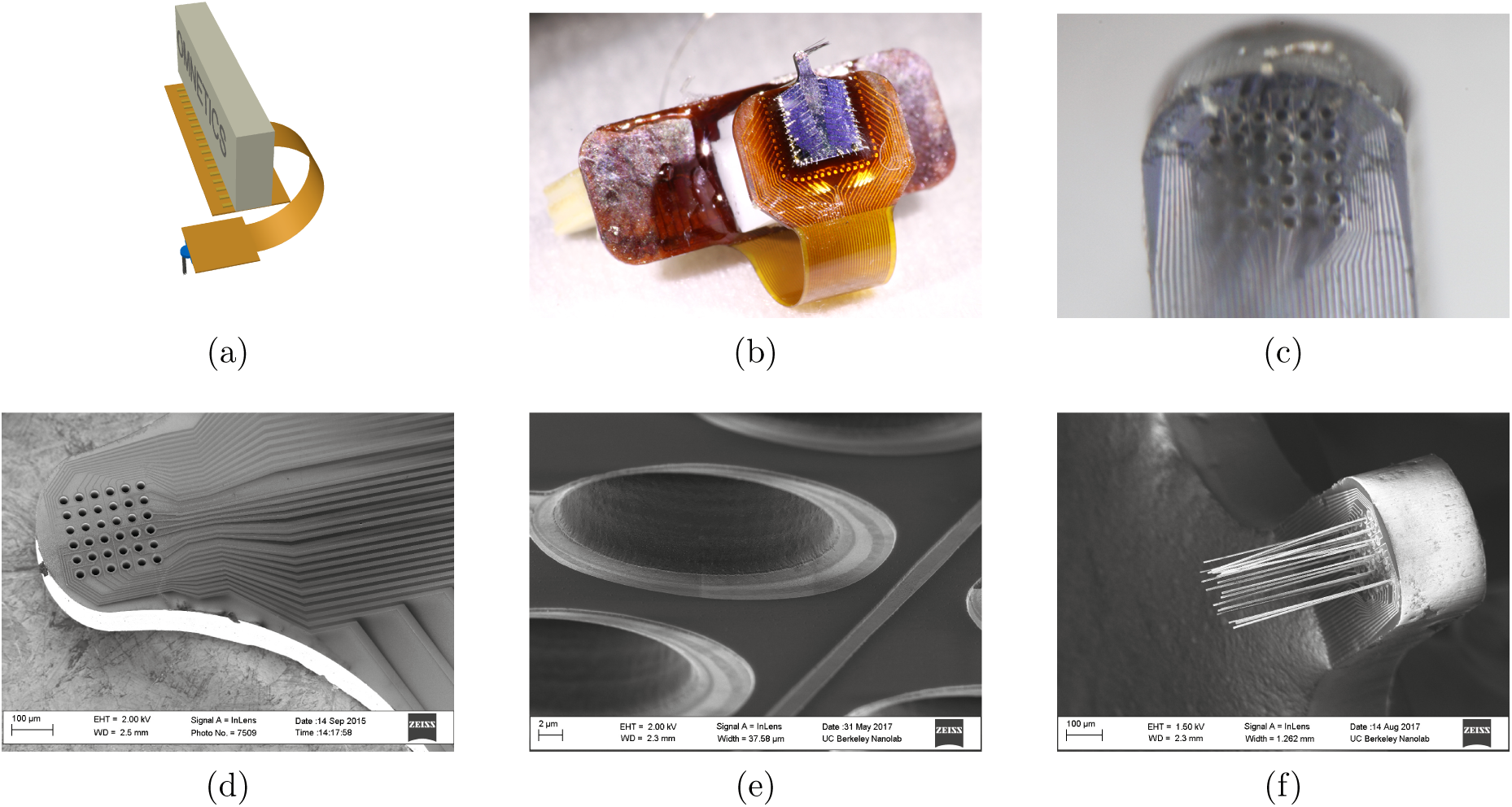
(a) Model of a complete assembled device showing the substrate (blue) with fibers (black) on the polyimide flex PCB (orange) with Omnetics header (white). (b) Photograph of an assembled recording array. (c) Close-up view of the head of a recording array with fibers (out of focus) extending upward out of plane. (d) SEM of substrate, and (e) a close-up on a portion of the head. (f) SEM of the head of a recording array with the carbon fibers clearly visible. Fibers in (b),(f) shortened for ease of imaging.

Substrate microfabrication yield was 60-70%, limited by the patterning of the 2 µm line, 4 µm space metallization between the vias. Surface tension effects due the topography, as well as inconsistencies in the thickness of the photoresist due to the manual dispense process, resulted in all metal traces on a given substrate either shorting or being completely removed in some regions of the wafer. Defective substrates were quickly identified by eye and removed, and with 912 substrates per wafer this did not present a significant limitation, particularly as it was the only yield-limiting step in the microfabrication process.

The assembly process resulted in a much higher yield, with greater than 90% of recording sites showing continuity (|*Z*| <10 MΩ) and greater than 80% showing impedances below 2 MΩ at 1 kHz across all devices. Yield during assembly is limited primarily by continuity through the silver ink, as discussed further in Section 4.

Impedance measurements for a representative device after exposing recording sites, after applying a breakdown voltage to the silver ink, and after electroplating the recording sites with PEDOT:PSS are summarized in Figure 4. Briefly, impedance consistently decreased by 2-10x upon applying a voltage across the electrode array, with those sites having higher initial impedance decreasing by a larger factor, and phase trending nearer to zero (more resistive). Impedance decreased further upon electroplating, typically approximately 1.5-5x, with negligible change in phase.

**Figure 4.**
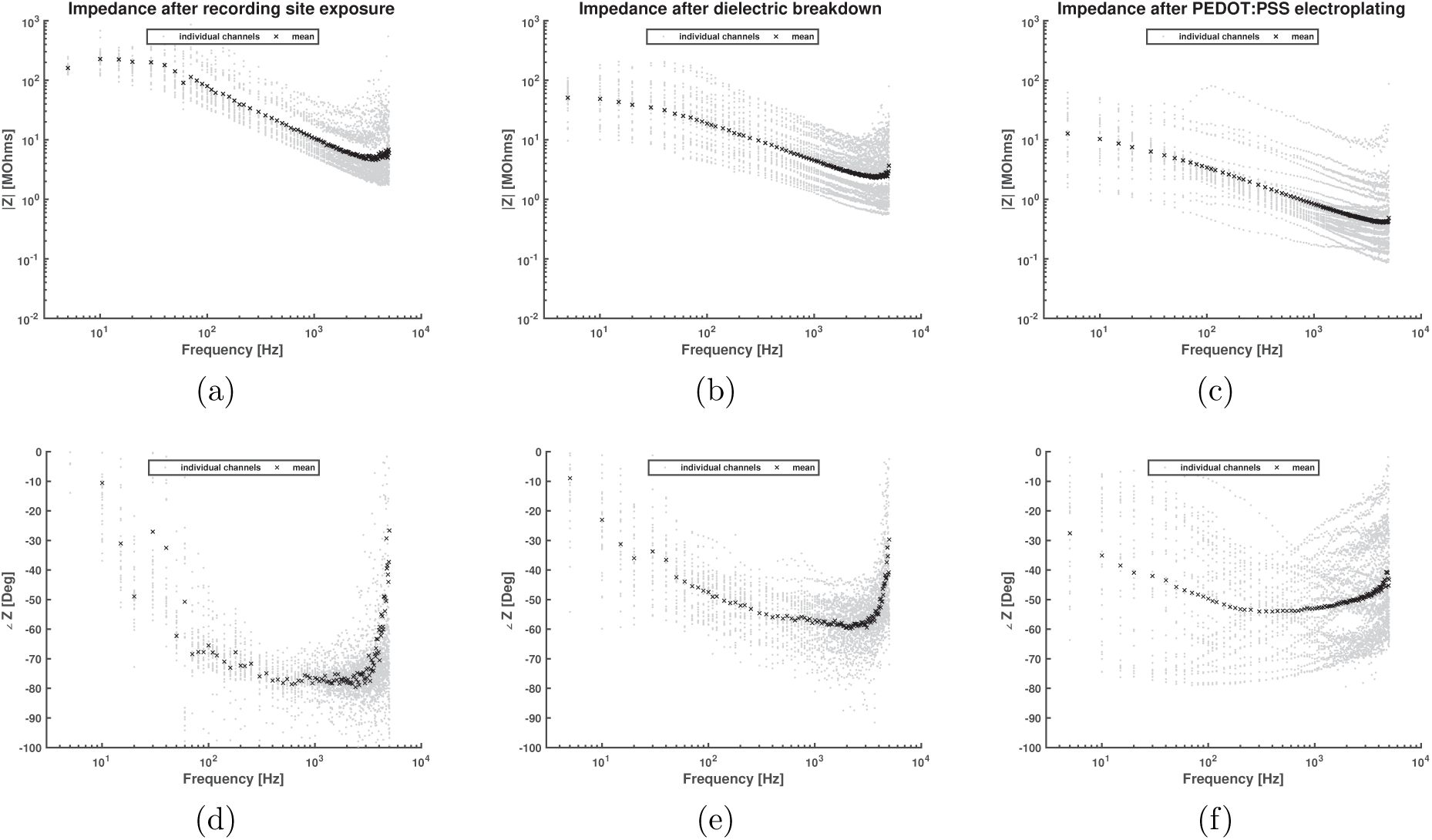
Impedance spectroscopy for electrodes on a typical device (a,d) immediately after recording sites are exposed, (b,e) after 18V is applied to break down residual dielectric in the silver ink, and (c,f) after electroplating the recording sites with PEDOT:PSS. The geometric mean and geometric standard deviation of the magnitude of impedances at 1 kHz are in (a) 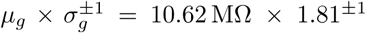, in (b) 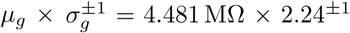, and in (c) 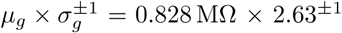.

To confirm that the silver ink dominated the electrode impedance before applying the breakdown voltage, impedance was measured between every pair of electrodes with recording sites in liquefied Field’s metal both before and after applying the breakdown potential. Similar impedance values measured in PBS and Field’s metal (Figure S3) suggest that the impedance before application of the breakdown voltage is dominated by the silver ink rather than by interface between the electrode and the surrounding medium. After applying the 18 V potential, 100 Hz impedance was observed to decrease by 2x in Field’s metal compared to PBS.

Thermal noise, which is a function of the electrode impedance and has a direct impact on the SNR of the recordings, was measured to be on average 8.4 µV over the 0.3-6 kHz band. The amplifier nominally contributes 1.3 µV (17.5 nV Hz^−0.5^) of noise uncorrelated to the electrode noise, and thus 8.3 µV (110 nV Hz^−0.5^) is attributable to the electrodes themselves. Spectral analysis confirmed that the measured noise was white; however, measurements taken on the Plexon recording system outside the Faraday cage showed greater electromagnetic interference. Spike-band noise density on the Plexon system is 450 nV Hz^−0.5^, approximately four times larger (41 µV over the 0.5-8.8 kHz band). Noise in the Plexon system’s 3-200 Hz band was 15 µV (1100 nV Hz^−0.5^). This increase in noise density at low frequencies is consistent with 60 Hz interference and its harmonics visible in spectral analysis (not shown).

Crosstalk is likewise an important consideration in the evaluation of a neural recording array, and can be estimated by the ratio of the impedance between any pair of electrodes while out of solution (open circuit) and in PBS. Figure 5 shows the mean impedance measured in air and in PBS, where the mean was taken over all 496 pairwise combinations of the 32 electrodes. A control experiment is also shown where no device was connected to the measurement apparatus, confirming that the measured impedance of the device in air is very similar both in magnitude and purely capacitive phase to the control, and that the extent of the crosstalk is below what can be measured with the current setup. The finite impedance is attributable to the non-zero CMOS off current in the multiplexer [35].

**Figure 5.**
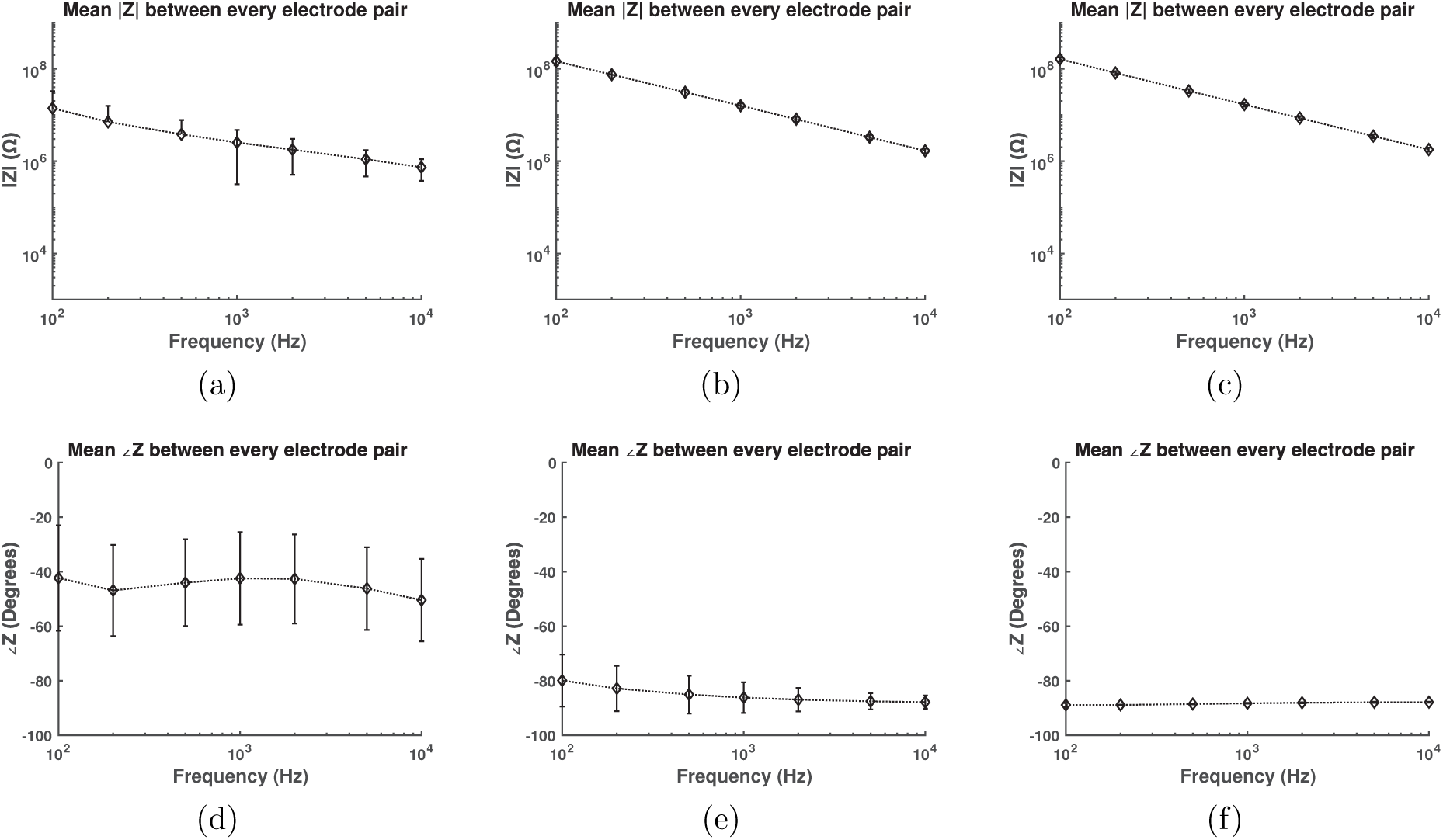
Average impedance (*µ* ± *σ*) between every pair of electrodes (a,d) in PBS, (b,e) in air, and (c,f) with no device connected as a control. The similar impedance values between the air and control cases suggest that the measured crosstalk is through the multiplexer rather than through the device itself, and thus that the crosstalk through the device is quite low.

## 3.2. Array penetration is strongly dependent upon fiber length but not upon angle

Testing the effect of fiber length and angle revealed that the angle of the fibers plays essentially no role with regards to the success of insertion within the range of angles tested, up to 4.5 degrees off-axis. This is presumed to be because local dimpling at the tip of each fiber effectively presents an orthogonal surface. This result suggests that while highly parallel fibers may have benefit for the distribution of recording sites, parallelism is not strictly required for successful insertion.

The length of the fiber, however, plays a strong role as expected from column buckling theory and as shown in [22]. Fibers shorter than 3.5 mm penetrated the 0.6% agar gel successfully on every attempt, but 3.5 mm fibers penetrated only with difficulty. Typically either multiple attempts were required, or some small lateral movement of the array was necessary while the fiber tips were in contact with the agar in order to coax the fibers to penetrate the surface. This set a practical upper bound of 2.5-3 mm for devices to be implanted *in vivo*, recognizing that agar is not a perfect model for cortical tissue.

## 3.3. Action potentials can be recorded in the CNS on multiple recording sites

The 32-channel array was used to measure spontaneous neural activity in M1, recording field potential on all channels and identifying well-isolated units on 20 channels. Figure 6 shows photos of the implanted array and summarizes key data validating the carbon fiber array for single- and multi-unit recording. The raster plot indicating unit timing and firing rate suggests that the majority of unit activity is likely produced by one neuron, with a limited number of other neurons producing the remainder of detected spike events. This is further supported by the high correlation observed among units and spatial proximity of the recording sites. Representative single-unit spikes with peak-to-peak voltage ranging from 52 µV to 115 µV are provided alongside a histogram of peak-to-peak voltages of detected units. Finally, field potential data from a single channel is shown in Figure S4.

**Figure 6.**
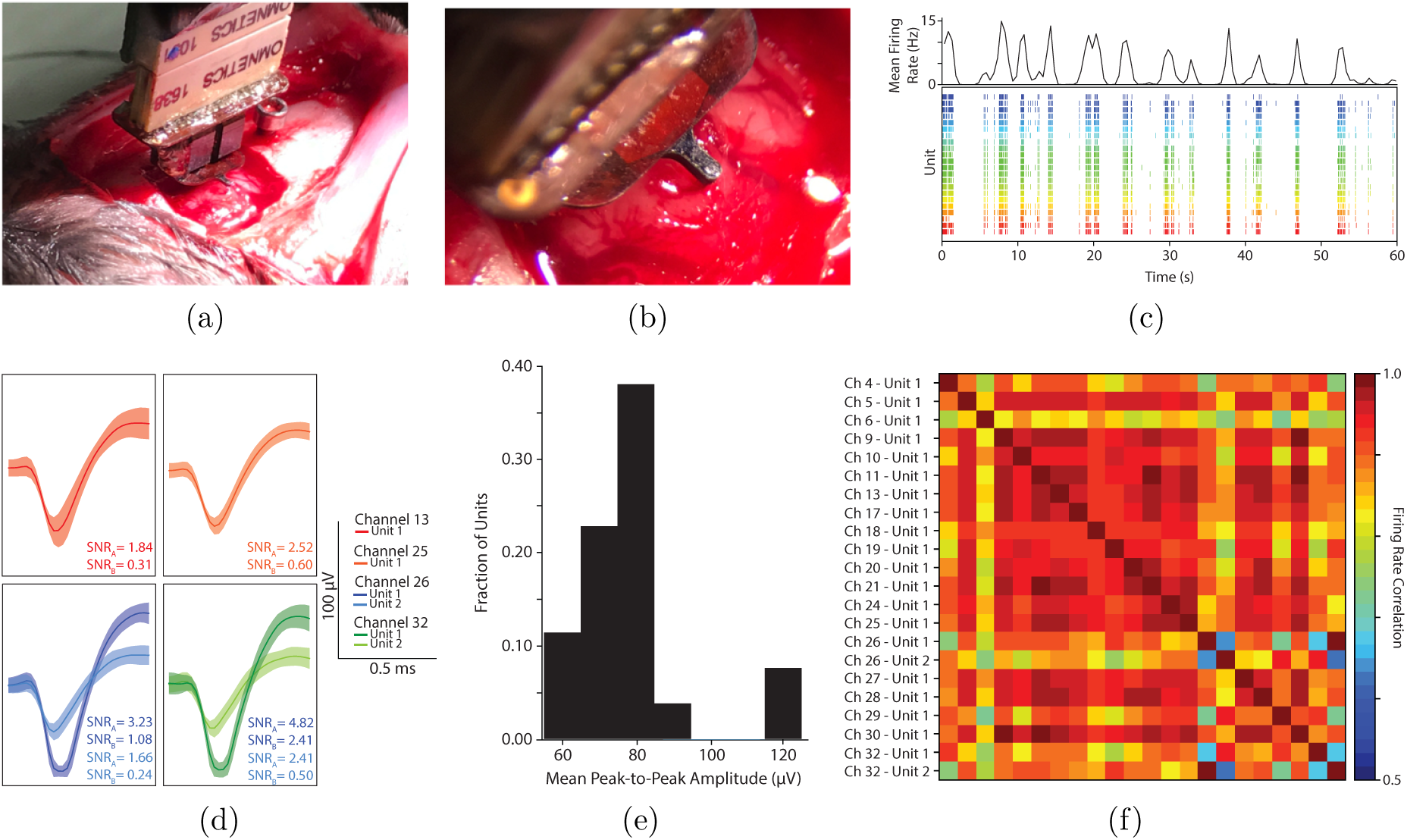
(a,b) Photographs of of a carbon fiber neural recording array implanted in M1. An enlarged craniotomy is shown for visibility, but in practice the craniotomy was approximately 1 mm in diameter. (c) Unit firing activity over time. The lower figure depicts a raster plot of the recorded units for 60 seconds, where each color represents spiking data recorded from a different unit. 22 units were recorded in total over 20 of the 32 channels. The upper figure displays the instantaneous firing rate averaged across all of the recorded units. (d) Representative unit waveforms recorded from M1. Mean unit waveforms from four channels are shown, with the shaded regions indicating the standard deviation of the corresponding unit activity. SNR values are given for each unit according to Methods A and B described in Section 3.3. (e) Distribution of peak-to-peak unit amplitudes. The amplitudes are grouped to into 10 µV bins to determine the distribution of waveform amplitudes. (f) Pearson correlation of firing rate across channels, with color indicating the strength of the correlation. Unit activity is binned into 500 ms segments over the duration of the recording and using to compute firing rates over time per unit. The unlabeled x-axis is identical to the labeled y-axis.

SNR is calculated for each recorded unit and tabulated in Table S2. Two methods are used, as there is no standardized method for computing the SNR of spikes given that they are nonperiodic signals. Method A takes the ratio of the peak-to-peak waveform voltage on a given channel to the RMS noise voltage. Method A is common in the literature [36–38]. Method B takes the square of the ratio of the RMS value of the spike waveform to the RMS noise voltage. While Method B resembles the traditional method of calculating SNR for a periodic signal, for a nonperiodic signal such as a spike waveform it is dependent upon the time window over which the spike RMS value is calculated. SNR values for Method A range from 0.85 to 4.8, and those for Method B range from 0.061 to 2.4.

## 4. Discussion

### 4.1. The microfabrication process is robust

The microfabrication process presented in Section 2 has been designed to be robust, and many of the design decisions reflect optimizations and trade-offs critical to the repeatability of the process. First among these decisions was the choice to use thin wafers. Inherent to DRIE is an aspect ratio limitation, such that there is a practical limit to the depth of an etched feature [39]. Even upon optimizing the etch process by increasing process pressure and bias power as a function of depth, the deepest 20 µm diameter cylindrical hole we could etch was 350-400 µm deep. Thinning a standard 525-675 µm thick wafer late in the microfabricaton process would compromise the backside insulating silica and present an opportunity for electrical shorting through the substrate; thus, beginning with a 280 µm thin wafer was deemed necessary.

Processing thin wafers comes with specific challenges, however. First, the wafers are outside the focal range of lithographic steppers, necessitating bonding the wafer to a 400 µm handle wafer as previously described. Second, because the thin wafer is more fragile than a typical wafer, standard temporary bonding methods present an elevated risk to the wafer, whether from trapped air bubbles cracking the wafer under vacuum or the debonding process itself requiring moderate flexion to separate the wafers. A water droplet provided the necessary bonding force for the gentle atmospheric environment of the lithographic stepper, and dicing tape carefully applied to the backside of a wafer allowed processing on low-temperature tools with vacuum chucks, namely photoresist coaters and developers. For plasma etching, in which neither water or dicing tape was suitable, we developed the aforementioned custom wafer bonding tool to enable uniform, void-free bonding of two wafers using a thin film of polyphenyl ether. Lastly, in addition to the aforementioned bonding challenges, the thin wafers were prone to developing dislocation defects in the oxidation furnace that resulted in significant warping at the edge. Because the wafers were no longer flat, additional care was necessary during bonding and cleaning to minimize pressure applied at the edges of the wafers that could result in hairline crack formation.

Hydrogen annealing was helpful following DRIE in order to smooth the sidewalls for two reasons. First, the sidewall scallops resulting from DRIE would shadow the sputtered aluminum deposition, resulting in a discontinuous film along the sidewall of the via. While the TiN film deposited by PEALD ensures coverage of the via sidewall, this additional coverage with aluminum reduces the resistance of the conductive film stack near the top of the via and doubly ensures continuity where it is most critical. Second, this annealing process results in rounding at the lip of the via, which makes it more energetically favorable for photoresist to enter the vias. This photoresist not only protects the vias and their sidewall metallization from the subsequent etch processes, but also greatly diminishes the streaking effect of spinning photoresist over deep topography.

To this latter point, spinning and patterning the photoresist near the vias presented a particular challenge, as the finest features in the process needed to be resolved immediately beside and between these aggressive topographical features. Because such topography is known to result in photoresist streaking during the spin process, and the dense square grid of holes presents an unfavorable energy landscape for photoresist coverage, the specific spin-coating process described above was critical. The dynamic dispense allows coverage of the holes without trapped air inside the vias creating thick bubbles of photoresist around the holes, while still avoiding surface energy-generated voids of photoresist in a close radius around the entire array of vias. The slow ramp rate and spin speed ensure that the photoresist doesn’t streak and is not removed from the vias, while still being fast enough to provide complete wafer coverage and uniformity of the photoresist film.

### 4.2. The properties of the isotropically conductive adhesive are critical for conductivity at small scales

Several aspects of the assembly procedure likewise warrant note for their subtle importance. Primary among these is the choice of isotropically conductive adhesive (ICA) for use in electrically and mechanically bonding the fibers and the vias. ICAs, and silver-filled epoxies in particular, are commonly used in microassembly of carbon fiber-based recording electrodes [20, 22, 24, 27], but our early experiments yielded inconsistent connectivity between the fiber and the conductive inner sidewall of the substrate.

Most ICAs operate by the formation of percolation networks, with the contact among many randomly arranged particles in the bulk material forming a network of conductive paths between the two relevant surfaces to be electrically (or thermally) connected. The ICA’s conductive particle (i.e. silver) content required to form a conductive network is function primarily of the average size, distribution, and shape of the particles in the ICA [40]. Size plays the most significant role, with a greater volumetric particle content, or fill, required relative to polymeric binder as the average particle size decreases.

In a constrained volume such as the vias on the array substrate, the largest particles in the distribution may be physically excluded. This has the effect both of reducing the average particle size and reducing the fill of the ICA. This latter effect can be significant, as large particles accounting for a negligible fraction of the total particle count can account for a substantial fraction of the total fill due to cubic scaling. One percent of particles by count in off-the-shelf silver epoxy formulations tested during development of this process, H20E and H20S (Epoxy Technology, Billerica, MA), may be as large as 45 and 20 µm, respectively, and the exclusion of these large particles is sufficient to decrease the epoxy fill below the conduction threshold on the majority of electrodes.

While it would initially seem straightforward to seek a silver epoxy formulation with a smaller average particle size or tighter distribution (AA-DUCT 24 was developed in part for this work and is quoted as having particles no larger than 2 µm), this correspondingly increases the required fill fraction, as noted above. An increase in the silver fill comes at the expense of the polymeric binder, which contracts as it cross-links during the cure to stress the silver particles and force them into intimate contact. With insufficient binder, this critical step in the process doesn’t occur and no conductive path will be formed, despite the high silver fill. Thus, a minimum amount of binder is also required to form a conductive network. Given that smaller particles require an increase in silver fill without decreasing the binder content, it becomes apparent there is a minimum particle size below which silver epoxies cannot form a percolation network, and indeed H20S (1-2 µm mean particle size) is near this lower limit.

Silver ink operates by a similar but subtly different mechanism from silver epoxies, in that there is a third key component in addition to the silver and the polymeric binder. Inks also contain a solvent, which vaporizes during cure to aid in effecting a volume loss to draw the particles into contact. As a result, less polymeric binder is required, and the effective silver fill after solvent evaporation can be higher than in an epoxy. Correspondingly, slightly smaller silver particles (Novacentrix HPS-030LV: 400-800 nm) can form a conductive percolation network.

The presence of a solvent comes at a cost, however; it begins to evaporate at room temperature, after 20-30 minutes a skin impenetrable by the carbon fibers forms on the surface. To prevent this skin formation and extend the working time of the silver ink, solvent evaporation must be inhibited with a cap layer that won’t interfere with the ink’s chemistry. After exploring unsuccessful options including adding low vapor pressure solvents atop the ink, the silver epoxy we had explored initially proved to be the best candidate. Using an epoxy cap layer additionally served to mechanically reinforce the joint, as silver epoxy is significantly stronger than silver ink.

### 4.3. The silver ink impedance can be further reduced

Despite optimizing the ICA formulation, the impedance contribution of the silver ink still dominated the overall impedance of most channels. To confirm, we tested the impedance of an electrode array in saline and compared against similar impedance measurements taken with the recording sites in Field’s metal, and found that the impedances were similar (Figure S3). This suggested that there was still a thin residual film of polymeric binder between silver particles in the ICA. Given the sub-micron size scale of silver particles, we hypothesized that the residual dielectric must be less than 100 nm in thickness, and breakdown of that dielectric would result in pyrolysis of the polymer and the formation of a graphitic short between silver particles. Typical values for polymer breakdown voltages range from 20-200 MV/m, so the expected breakdown potential was expected to be less than 20 V. Indeed this is what we found, with an application of −18 V DC (per Section 2) proving sufficient to reliably reduce the impedance by 0.5-1.5 orders of magnitude. The formerly highest impedance recording sites were reduced most significantly, with the overall effect that the variability in electrode impedance within each device was substantially reduced following this dielectric breakdown treatment. Below −18 V, the submerged portion of the carbon fibers could be destructively oxidized due to the energy provided by the large potential. With the electrodes reversed, such that electrolysis produced oxygen at the carbon fibers rather than hydrogen, the voltage at which oxidation occurred was significantly lower.

### 4.4. Mitigating electrostatic interactions during assembly is necessary for fibers to remain parallel

The fibers acquire electrostatic charge during assembly, causing the fibers to repel each other and deflect outward. If the silver ink and epoxy are cured while the fibers are electrostatically splayed, they will maintain some of that divergence even if later discharged. Steps can be taken to minimize electrostatic charging of the fibers during assembly, including increasing the ambient humidity and using a neutralizing ion generator, and indeed this has merit in mitigating complicating interactions among fibers or between fibers and other objects during assembly, but it is impossible to prevent the accumulation of some charge. Thus, it is critical that the fibers be discharged until after the ICAs are cured. This is most conveniently achieved with the application of the aluminum strip short circuiting all channels to ground during assembly.

### 4.5. Embedding temperature, compound, and blade choice are critical to cleanly exposing recording sites

In exposing the recording sites, several parameters were experimentally varied to qualitatively achieve the cleanest possible cut. After each trial, the fibers were examined under an optical microscope and/or SEM to assess the length of fiber removed, the number of fibers cut, the occurrence of incompletely severed parylene insulation, and stretching of the parylene insulation beyond or over the tip of the fiber. The carbon fiber itself was trivial to cut (or break) through; parylene is more challenging. Temperature, embedding compound, and cryostat blade choice each play a significant role in the quality of the cut through the parylene-coated carbon fiber.

Given the above, the goal is to cut through the majority of parylene-coated fibers on every pass with minimal inelastic deformation of the parylene. Thus, the ideal embedding compound has similar mechanical properties to the parylene insulation. Because the hardness and compliance of embedding compounds is a strong function of temperature, and embedding compounds are generally much softer and more compliant than parylene, lower temperatures are favorable. We tested the standard Tissue-Tek OCT (optimal cutting temperature) compound at −26 °C, −38 °C, and −55 °C, and found that the cuts were consistently of the highest quality at −55 °C. We observed greater elongation of the parylene at higher temperatures, and at higher temperatures each fiber was cut only every second or third 10 µm pass of the cryostat.

Based on the observation that temperature and thus hardness was critical, we also tried an embedding compound designed for low-temperature use and water ice. The low-temperature embedding compound was actually softer than the standard embedding compound at a given temperature, having been designed for sectioning lipid-rich tissue at −40 °C. The water ice, which was considerably harder than both polyvinyl alcohol and polyethylene glycol-based embedding compounds, was unacceptably brittle and resulted in cuts of widely variable thicknesses as the blade struggled to engage such a thin layer of ice.

Lastly, the exact blade type matters. Infinity, Gold, Extremus, and Diamond blades were purchased from C.L. Sturkey, Inc., Lebanon, PA. Infinity blades had no advertised ceramic coating, but were ground with three bevels; gold blades were coated in a titanium nitride thin film; diamond blades were coated in an amorphous diamond thin film; and extremus blades were coated in an unknown film, advertised as being well suited to a wide variety of cutting conditions. Upon examining fibers cut by each blade under otherwise ideal conditions under an SEM, the “Gold” blades consistently produced the cleanest cuts, closely followed by Infinity and Extremus with more frequent elongation of the parylene. The Diamond blades performed poorly for this application, resulting in many broken fibers still attached by incompletely severed parylene.

### 4.6. This design presents a path toward scalability

For a device to be practically scalable, two requirements must be met. First, any steps that scale with the number of electrodes must be automated. Second, basic electronics (amplification, multiplexing, and digitization) should be integrated into the device itself to minimize the number of connections between the recording array and headstage. The only step in device creation that scales with the number of recording sites is the threading of fibers, which we have shown can be automated [28]. Integrating electronics onto or into the substrate is also possible. While post-processing CMOS wafers with our MEMS fabrication processes would be possible with some modification of processing to eliminate high-temperature steps, a more straightforward approach would involve leveraging the silver ink to electrically connect the array substrate to an array of pads (analogous to flip-chip bond pads) on a CMOS die with the electronics. This would add only the assembly step of aligning the array substrate to the CMOS die, and would dramatically simplify the substrate microfabrication and improve yield as metallization would no longer be necessary.

## 5. Conclusions

We have successfully demonstrated a high-density 32-channel carbon fiber neural recording array with 5.2 µm electrodes at 38 µm pitch. The silicon microfabrication process presented is scalable to both a large number of devices and a large number of recording sites, and is robust and repeatable. Likewise, the assembly process has only one serial (per-fiber) step, and that step may be automated, meaning that the process is again scalable to a large number of recording sites. This assembly process has been optimized for high yield and minimum recording impedance; characterization data show sub-megaohm impedance values well suited to single-unit recording and negligible crosstalk among channels. *In vivo* experiments have confirmed the devices’ capability to record single- and multi-unit action potentials on a large number of recording sites in addition to local field potential.

The demonstration of such a device is a step forward in acute intracortical recording, as microwire arrays and their Utah variants remain the workhorses of high-channel-count neural recording in the central nervous system, particularly for brain-machine interface applications [41, 42]. This device presents a feasible roadmap toward scalability both with regards to fabrication techniques as well as the potential for CMOS integration, while continuing to minimize volumetric displacement of neural tissue by minimizing the electrode size.

## Acknowledgments

This work was supported by DARPA VAPR HR0011-14-2-0001, NIH R21 EY027570, and DARPA ELM DE-AC02-05CH11231. We are grateful to Camilo Diaz-Botia for providing the MATLAB scripts for EIS; Amy Liao, Monica Lin, and Sarah Swisher for the multiplexer hardware and software for pairwise impedance measurement; and Steve Ruzin and Denise Schichnes of the College of Natural Resources (CNR) Biological Imaging Facility for use of the cryostat. In addition, we thank Bill Flounders and the staff of the Marvell Nanofabrication Laboratory, the Berkeley Sensor and Actuator Center (BSAC), the Ubiquitous Swarm Lab at UC Berkeley, and the Center for Neural Engineering and Prostheses (CNEP) for their continued support. Michel Maharbiz is a Chan Zuckerberg Biohub investigator.

